# Spike Protein of SARS-CoV-2 Activates Macrophages and Contributes to Induction of Acute Lung Inflammations in Mice

**DOI:** 10.1101/2020.12.07.414706

**Authors:** Xiaoling Cao, Yan Tian, Vi Nguyen, Yuping Zhang, Chao Gao, Rong Yin, Wayne Carver, Daping Fan, Helmut Albrecht, Taixing Cui, Wenbin Tan

**Author notes:** contributed equally (Xiaoling Cao, Yan Tian, Vi Nguyen). **Correspondence:** Wenbin Tan, Department of Cell Biology and Anatomy, School of Medicine, and Biomedical Engineering Program, College of Engineering and Computing, University of South Carolina, Columbia, South Carolina, 29209, USA. **Disclaimer** Any views expressed here represent personal opinion and do not necessarily reflect those of the U.S. Department of Health and Human Services or the United States federal government. **Prior Publication:** None of the material in this manuscript has been published or is under consideration for publication elsewhere, including the Internet.

## Abstract

**Background:** Coronavirus disease 2019 (COVID-19) patients exhibit multiple organ malfunctions with a primary manifestation of acute and diffuse lung injuries. The Spike protein of severe acute respiratory syndrome coronavirus 2 (SARS-CoV-2) is crucial to mediate viral entry into host cells; however, whether it can be cellularly pathogenic and contribute to pulmonary hyper-inflammations in COVID-19 is not well known.

**Methods and Findings:** In this study, we developed a Spike protein-pseudotyped (Spp) lentivirus with the proper tropism of SARS-CoV-2 Spike protein on the surface and tracked down the fate of Spp in wild type C57BL/6J mice receiving intravenous injection of the virus. A lentivirus with vesicular stomatitis virus glycoprotein (VSV-G) was used as the control. Two hours post-infection (hpi), Spp showed more than 27-75 times more viral burden in the lungs than other organs; it also exhibited about 3-5 times more viral burden than VSV-G lentivirus in the lungs, liver, kidney and spleen. Acute pneumonia was evident in animals 24 hpi. Spp lentivirus was mainly found in LDLR^+^ macrophages and pneumocytes in the lungs, but not in MARC1^+^ macrophages. IL6, IL10, CD80 and PPAR-γ were quickly upregulated in response to infection of Spp lentivirus in the lungs *in vivo* as well as in macrophage-like RAW264.7 cells *in vitro*. We further confirmed that forced expression of the Spike protein in RAW264.7 cells could significantly increase the mRNA levels of the same panel of inflammatory factors.

**Conclusions:** Our results demonstrate that the Spike protein of SARS-CoV-2 alone can induce cellular pathology, e.g. activating macrophages and contributing to induction of acute inflammatory responses.

## Introduction

Coronavirus disease 19 (COVID-19) has become a significant threat to global health. So far, more than 64 million infections and 1.5 million victims have been reported worldwide including 191 countries and regions. The US alone has registered more than 14 million cases and 271 thousand deaths as of date December 4, 2020 [1]. Severe acute respiratory syndrome coronavirus 2 (SARS-CoV-2) is the causative organism for COVID-19 [2]. SARS-CoV-2, a positive-sense single-stranded RNA virus, is the newest and seventh known coronavirus that is capable of infecting humans [2, 3]. The SARS-CoV2 Spike (S) protein can be cleaved by Furin to produce subunits S1 and S2 to form a trimer which can mediate viral entry into host cells via surface angiotensin converting enzyme 2 (ACE2) [4]. Host Transmembrane Serine Protease 2 (TMPRSS2) is also capable of promoting SARS-CoV-2 entry of target cells by cleavage of the S2’ site in the S2 subunit [5, 6]. ACE2 and TMPRSS2 have been found to co-express in lung type II pneumocytes, ileal absorptive enterocytes, and nasal goblet secretory cells [7], which are thought to be host determinants for viral infection in the initial stage. Recently, Neuropillin (NRP) 1 has been identified as the second host factor to facilitate SARS-CoV-2 entry of target cells which appears to be ACE2-independent since a different binding motif is used [8–10].

COVID-19 patients can be asymptomatic or symptomatic. The mortality rate of the COVID-19 varies in different geographic locations and patient populations [1]. Patients with metabolic-associated preconditions such as hypertension, cardiovascular disorders (CVD), obesity and diabetes mellitus (DM) are experienced to develop more severe symptoms [11]. COVID-19 patients also showed decreases in serum lipid levels [12, 13]. SARS-CoV-2-induced hyper-inflammation in the lungs is considered to cause the disease progression. The molecular mechanisms of COVID-19 pathogenesis have just begun to be elucidated. In this study, we aim to investigate whether the S protein of SARS-CoV-2 interacts with macrophages and induce acute lung inflammations *in vivo* using a S protein-pseudotyped (Spp) lentivirus.

## Methods

### Materials

Phoenix cells and Dulbecco’s Modified Eagle’s medium (DMEM) were purchased from ATCC (Manassas, VA, USA). Lentiviral vector pLV-mCherry and vesicular stomatitis virus glycoprotein (VSV-G) expression vector pMD2.G were obtained from Addgene (Watertown, MA, USA). Coding sequence of SARS-CoV-2 S gene (GenBank: QHU36824.1) fusion with a c-terminal His tag was synthesized *in vitro* (Genscript, Piscataway, NJ, USA) after codon optimization for expression in human cells. The sequence was cloned into a pcDNA3.1 vector to obtain pcDNA-Spike. Anti-Spike S1 subunit, anti-low density lipoprotein receptor (LDLR), anti-mannose receptor C-type 1 (MRC1), anti-CD68 and anti-human immunodeficiency viruses (HIV)-1 p24 antibodies were obtained from Novus Biologicals (Littleton, CO, USA). The anti-His tag antibodies were obtained from Proteintech (Rosemont, IL, USA) and Thermo Fisher (Waltham, MA, USA). Primers were synthesized by IDT (Coralville, IA, USA) and primer sequences are listed in supplementary table 1. RNA extraction kit was obtained from Zymo Research (Irvine, CA, USA). The RT kit was obtained from Takara Bio USA (Mountain View, CA, USA). The SYBR green master mix was from BioRad (Hercules, CA, USA). Wild type C57BL/6J mice were purchased from the Jackson Laboratory (Bar Harbor, ME, USA).

### Generation of pseudotyped lentivirus

SARS-CoV-2 S gene (GenBank: QHU36824.1) fusion with a c-terminal 12xHis tag was synthesized and cloned into a pcDNA3.1 vector. Phoenix cells were grown in DMEM containing 10% FBS and co-transfected by pLV-mCherry and pcDNA-Spike or pMD2.G vector using a calcium phosphate kit (ThermoFisher, Waltham, MA, USA). The supernatant with produced virus (Spp or VSV-G lentivirus) was harvested 72-hours post transfection, clarified by centrifuging at 5000 g for 15 min followed by filtration of the supernatant through a 0.45 μm filter disk. The virus was collected by an ultracentrifugation at 24,000 rpm for 2 hours (hrs) using Beckman SW41 rotor. The viral pellets were resuspended by cold PBS buffer and stored at −80 °C before use. The viral particle number was determined using a real time RT-PCR assay to quantify the RNA copies of mCherry.

### Intravenous viral administration in vivo

The animal protocol was approved by the University of South Carolina IACUC committee. Male wild type C57BL/6J mice (5-6 weeks old) were intravenously administered 100 μl of Spp or VSV-G lentivirus (8×10^8^ of viral particles) via tail venous or retro-orbital injection. The animals were sacrificed at 2 or 24 hrs post-infection (hpi) and perfused by 50 ml PBS per mouse. The tissues including lungs, heart, liver, kidney, aorta, and spleen were collected. One part of tissue was used for RNA extraction followed by a real time RT-PCR to determine the number of viral particles in each tissue. The other part was fixed, embedded and used for histology and immunohistochemistry.

### RAW264.7 cell culture, viral uptake and electroporation

Macrophage-like RAW264.7 (RAW) cells (*ATCC*^®^ TIB-71™) were cultured in DMEM (10% FBS) medium. The cells were changed into 2% FBS DMEM medium for overnight prior to viral uptake assay. Spp or VSV-G lentivirus were added into RAW cells (4.8×10^7^ particles per well) in 12 well-plate with 90% confluence and incubated for 2 or 16 hours. Following treatment, the cells were washed with PBS three times and RNA was extracted using an RNA extraction kit (Zymo, Irvine, CA, USA). In a parallel experiment, RAW cells (5×10^6^) were electroporated with pcDNA3.1, EGFP-N2 or pcDNA-Spike plasmids (10 μg) using the following parameters: 2 mm gap cuvette, 250 ul sample volume and 120V (BTX Harvard Bioscience, Inc., Holliston, MA, USA). The cells were harvested 48 hrs post-electroporation for analysis.

### Real time RT-PCR and immunohistochemistry assay

To generate cDNA, 1.0~5 μg of total RNA was reverse-transcribed in a 20-μl reaction containing 1x RT buffer (Clontech, Mountain View, CA, USA), 0.5 mM dNTPs, 0.5 μg of oligo (dT) 15-mer primer, 20 units of RNasin, and 5 units of SMART Moloney murine leukemia virus reverse transcriptase (Takara Bio, Mountain View, CA, USA). The RT reaction was carried out at 42°C for 2 hrs. Seven house-keeping genes were screened for the normalization controls: GAPDH, Rps18, Ppia, Nono, Rpp30, Alas2, and β-actin. We found that Rps18 and Nono showed much more stable expression levels in tissues crossing various samples (data not shown); both Rps18 and Nono were then used as controls to normalize the amplification data. Expression levels of a panel of 23 inflammatory genes (Supplementary Table 1) were determined using real time RT-PCR. The reaction for the multiplex real time PCRs contained 1× SYBR Green qPCR Master Mix (Bio-Rad, Hercules, CA, USA), 10 ng of each template, and 10 pmol of each specific primer in a 25-μl total volume in a 96-well format. Each reaction was performed in duplicate under identical conditions. The PCR conditions were one cycle at 95°C for 2 min followed by 45 cycles of 15 s at 95°C and 60 s at 60°C. Relative quantification of the real time PCR was based upon the amplification efficiency of the target and reference genes and the cycle number at which fluorescence crossed a prescribed background level, cycle threshold (*Ct*).

For immunoblot assay, cell lysates were extracted from RAW cells using RIPA lysis buffer (Santa Cruz Biotech., Inc., Dallas, TX, USA). Proteins were separated by SDS-PAGE and transferred onto PVDF membranes. Anti-Spike S1 subunit, anti-HIV-1 p24 antibody, or anti-His antibodies were used to detect the expression of viral proteins and followed by HRP-labeled secondary antibodies. Images were acquired using a Bio-Rad Gel Imaging System (Hercules, CA, USA).

Lung tissues from mice were fixed in 10% buffered formalin (Fisher Scientific, Pittsburgh, PA, USA) and embedded in paraffin. Approximately 6 μm thick sections were cut and collected. The sections were blocked with 5% donkey serum and then incubated in a humidified chamber overnight at 4°C with primary antibodies. Sections were rinsed and incubated with fluorescent conjugated secondary antibodies for 2 hours at room temperature. Images were acquired using ImageXpress Pico System (Molecular Device, San Jose, CA, USA) or confocal microscopy system (Carl Zeiss AG, Oberkochen, Germany).

### Statistical analyses

All statistical analyses were performed in Origin 2019. The paired *t* test or one-way ANOVA was used for two groups or multiple comparisons test, respectively. The data was presented as “mean ± s.d.” and *p* < 0.05 was considered as significant.

## Result

### Tissue distributions of Spp in infected mice

We generated Spp lentivirus from Phoenix cells. Spike protein was shown to be cleaved into S1 and S2 subunits assembled on the Spp lentivirus that were produced in Phoenix cells by immunoblot analysis (Fig 1A). No evidence of full length, non-cleaved Spike protein was detected in the Spp lentivirus (Fig 1A). A lentivirus using a helper vector expressing VSV-G was obtained as a control, referred to as VSV-G lentivirus. Gag-p24 is the capsid core shell protein in both Spp and VSV-G lentiviruses which was shown in both Spp and VSV-g lentiviruses (Fig 1A).

**Fig 1.**
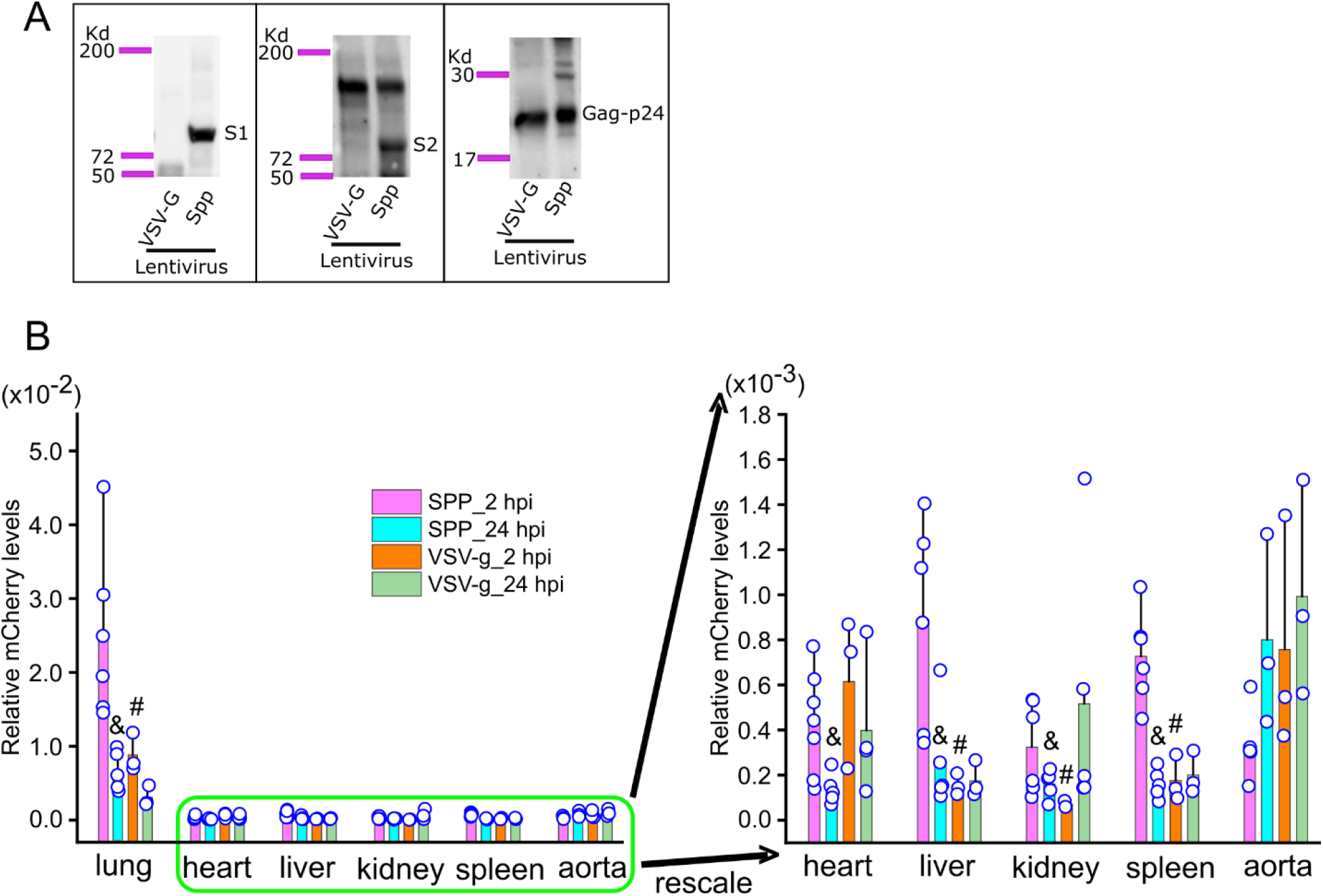
Systemic dissemination of Spike protein-pseudotyped (Spp) and vesicular stomatitis virus glycoprotein (VSV-G) lentiviruses in mice. (A) Detection of S1 and S2 subunits in Spp lentivirus by Western blot using a specific antibody against the S1 subunit and an anti-His antibody recognizing the S2 subunit. Both Spp and VSV-G lentiviruses have a gag-p24 protein. (B) Spp shows a predominant distribution in the lungs after being intravenously administrated. &, *p*<0.05, Spp viral burden as comparison with VSV-g in the same tissue at 2 hours post-infection (hpi); #, *p*<0.05, Spp viral burden at 24 hpi as comparison with 2 hpi in the same tissue.

Spp or VSV-G lentivirus (8×10^8^ particles) was intravenously injected into C57BL/6J mice. The animals were sacrificed at 2 or 24 hpi and various tissues were collected for determining the viral load. Both Spp and VSV-G had highest vial loads in the lungs at 2 hpi; Spp showed a factor of 27, 33, 55, 71 and 74 times the viral loads in the lungs compared to the liver, spleen, heart, aorta and kidney (*p*<0.05, n=3~7 mice); Spp also showed a factor of 2.8, 4.1, 4.5 and 5.7 times viral loads compared to VSV-G lentivirus in the lungs, spleen, kidney and liver, respectively (*p*<0.05, n=3~7 mice) (Fig 1B). At 24 hpi, the viral loads for Spp decreased significantly in the lungs, heart, liver, kidney and spleen (*p*<0.05, n=3~7 mice) (Fig 1B).

### Pathological features of pneumonia in Spp-infected mice

We then asked whether the mice treated with Spp virus acquired pneumonia. There were no evident histological changes in the lungs 2 hpi in both Spp and VSV-G groups. At 24 hpi with Spp lentivirus, many pathological changes in the lungs were evident, including multifocal lesions, inflammatory cell infiltrations, thickened alveolar walls, peri-vascular and peri-bronchial infiltrations and fibroplasia with exudation of fibrin and proteins (Fig 2). We observed only mild inflammations such as mildly thickened alveolar walls in the lungs in the VSV-G group at 24 hpi (Fig 2). Taken together, Spp but not VSV-G lentivirus could induce acute and diffuse pneumonia in the mouse lungs with very similar pathological manifestations observed in severe COVID-19 patients. These data also suggested that the S protein of SARS-CoV-2 played an important role in the development of acute pneumonia.

**Fig 2.**
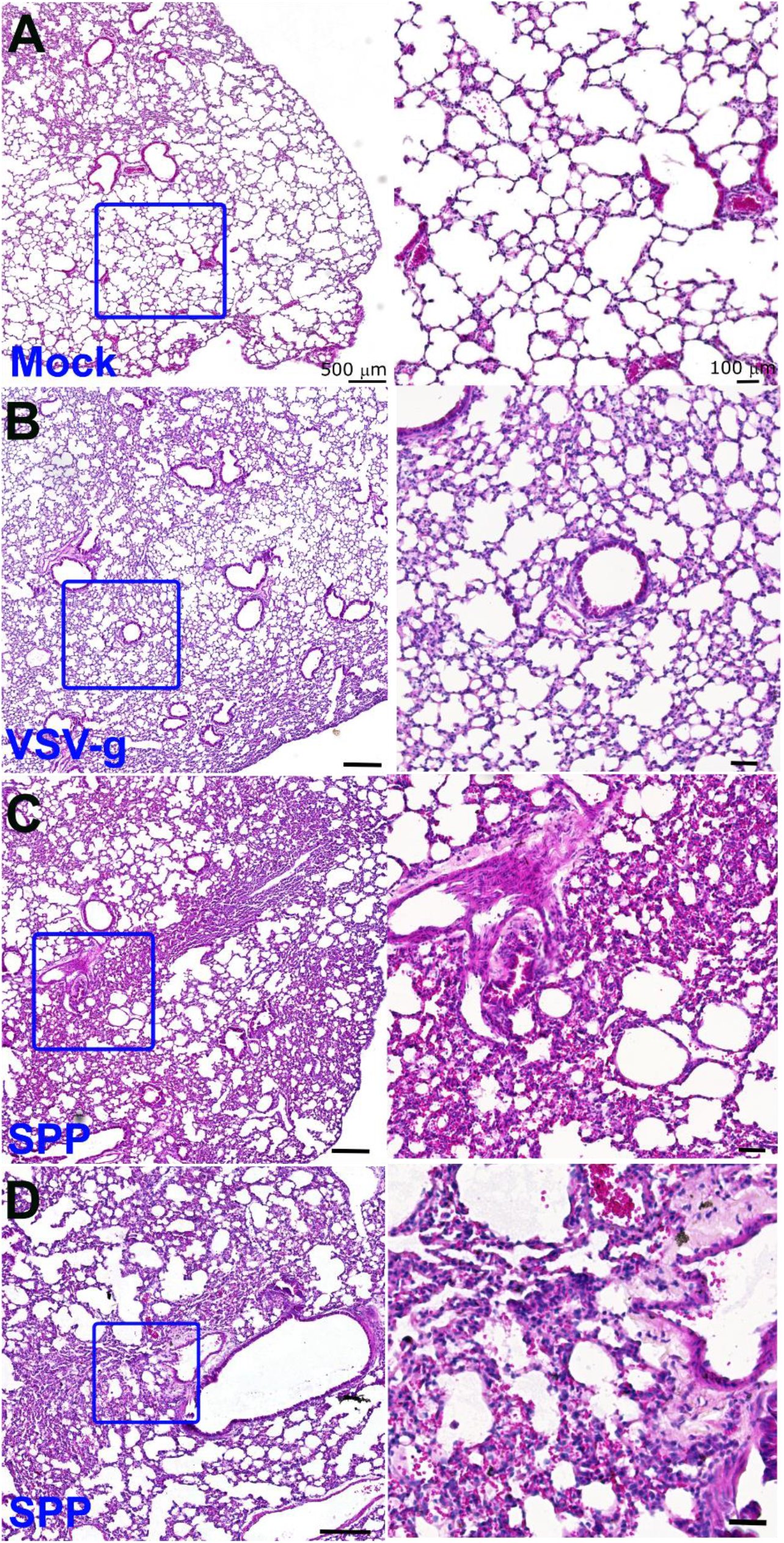
Mice acquires acute pneumonia 24 hours post-infection (hpi) of Spike protein-pseudotyped (Spp) lentivirus. Histological analysis of lungs in control mice (A), mice being administrated with lentivirus carrying vesicular stomatitis virus glycoprotein (VSV-G) (B), or Spp lentivirus (C and D) shows acute and diffuse inflammatory responses in the lungs after 24 hpi of Spp. Right panel is the magnification of boxed area in the corresponding left panel. Scale bar: 500 μm (left panel) and 100 μm (right panel).

### Cellular colocalization of Spp lentivirus in the lungs

We next examined cellular distribution of Spp lentivirus in the lungs. Spp viral antigen was detected by an anti-His antibody. LDLr is expressed in type II alveolar epithelial cells and macrophages in the lungs [14, 15]. The majority of cells that demonstrated Spp lentivirus-uptake (His^+^) (82.3%±11.4%) were LDLr^+^ cells, while 84.4%±14.9% of LDLr^+^ cells showed uptake of Spp lentivirus (Fig 3). We then examined the types of macrophages with uptake of Spp lentivirus in the lungs using macrophage markers CD68 and MRC1. We found 10%±4.4% of cells with evidence of Spp lentivirus uptake were CD68^+^ macrophages, while 38.3%±19.3% of CD68^+^ macrophages showed uptake of Spp lentivirus (Fig 3). However, we could find little evidence of MRC1^+^ macrophages that had uptake of Spp lentivirus (Fig 3).

**Fig 3.**
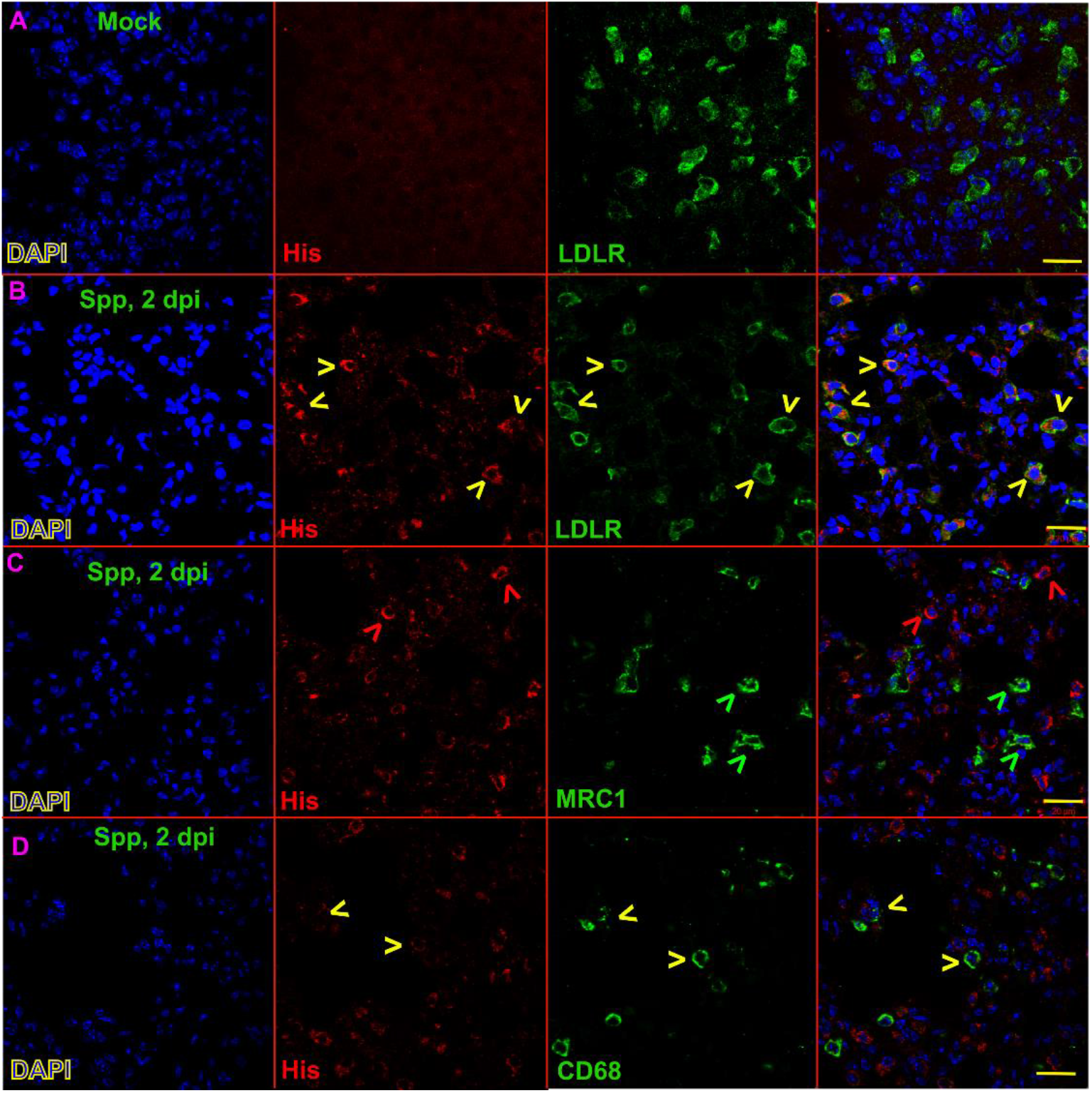
Cellular colocalizations of Spike protein-pseudotyped (Spp) lentivirus in the lungs 2 hours post-infection (hpi). An anti-his tag antibody is used to recognize S-fusion protein in Spp. Yellow arrowhead indicates cells positive for both His-tag and corresponding lung markers. Red or green arrowhead indicate cells positive for His tag or corresponding lung markers only, respectively. Scale bar: 20 μm.

### Dysregulation of inflammatory cytokines in the lungs in Spp-infected mice

We next attempted to assess which inflammatory factors might be induced by Spp lentivirus. We examined expression levels of a panel of 23 genes that are representative inflammatory markers in macrophages (Supplementary Table 1). Two hours post viral administration, the mRNA levels of IL 6, IL10, CD80 and PPAR-γ showed a significant and rapid increase in the lungs in Spp-infected mice but not VSV-g-infected mice as compared with untreated control mice (Fig 4). The levels of TNF-α showed an increase in mice both 2 and 24 hpi with Spp and VSV-g as compared with untreated control mice (Fig 4). TGF-β showed a significant increase in the lungs of Spp-infected mice only at 24 hpi as compared with untreated control mice and mice at 2 hpi (Fig 4). We did not find significant changes in expression levels of other inflammatory markers that we examined (data not shown). These data suggested that IL 6, IL10, CD80 and PPAR-γ were among those factors induced in a rapid response to the Spp infection.

**Fig 4.**
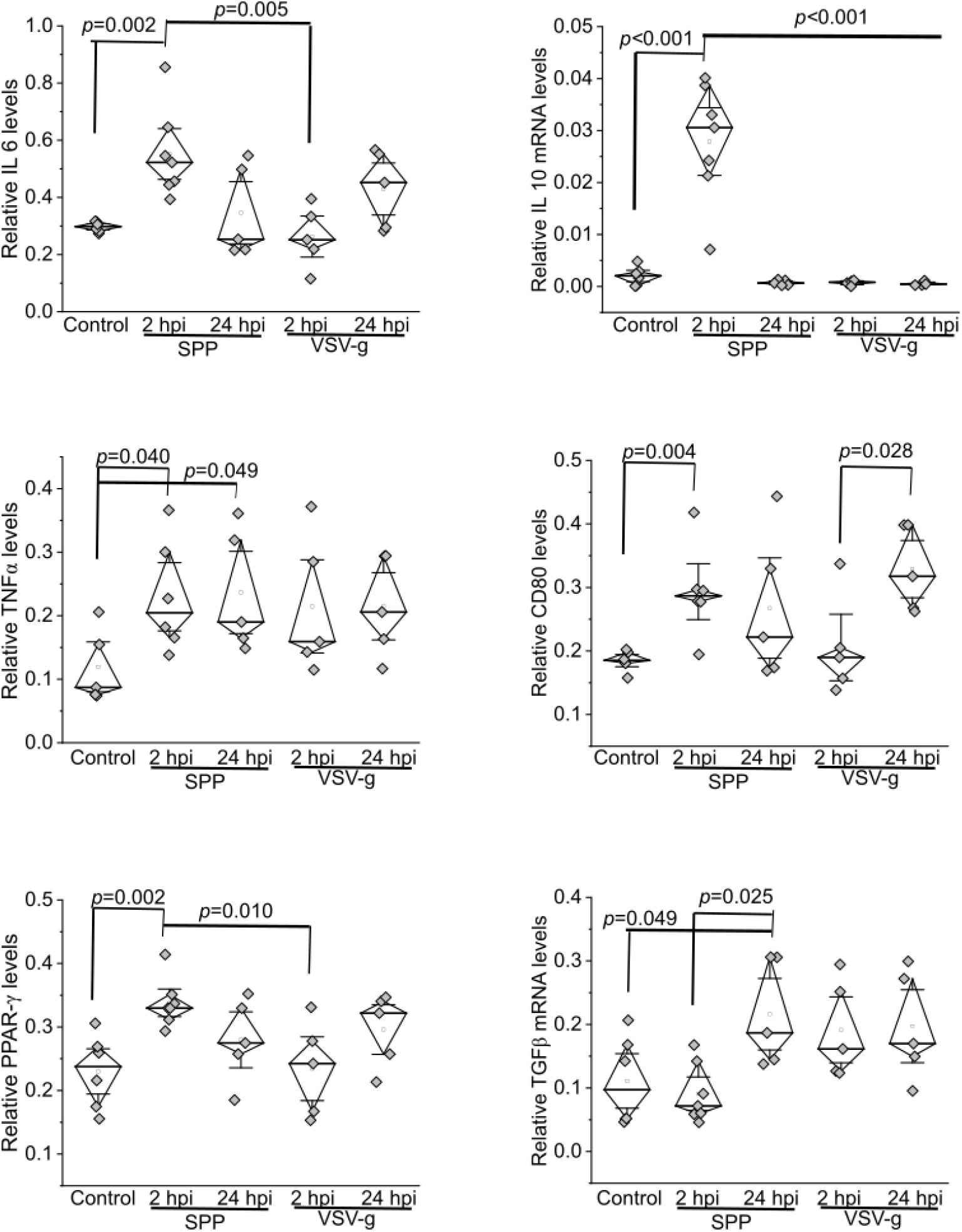
Upregulation of inflammatory factors in the lungs of mice after administration of Spike protein-pseudotyped (Spp) lentivirus or lentivirus carrying vesicular stomatitis virus glycoprotein (VSV-G). The mRNA level of each target gene is normalized to Rps18 levels.

### Dysregulation of inflammatory cytokines in RAW cells caused by the Spike protein

In order to investigate the potential immunomodulatory function induced by the S protein, we infected RAW cells using Spp or VSV-g lentivirus. The mRNA levels of IL 6, IL10, CD80 and PPAR-γ were significantly higher in the Spp-infected group at 2 hpi than untreated controls and VSV-G-infected group at 2 hpi (Fig 5). Their mRNA levels were continuously increased in the Spp-infected group at 16 hpi than 2 hpi (Fig 5).

**Fig 5.**
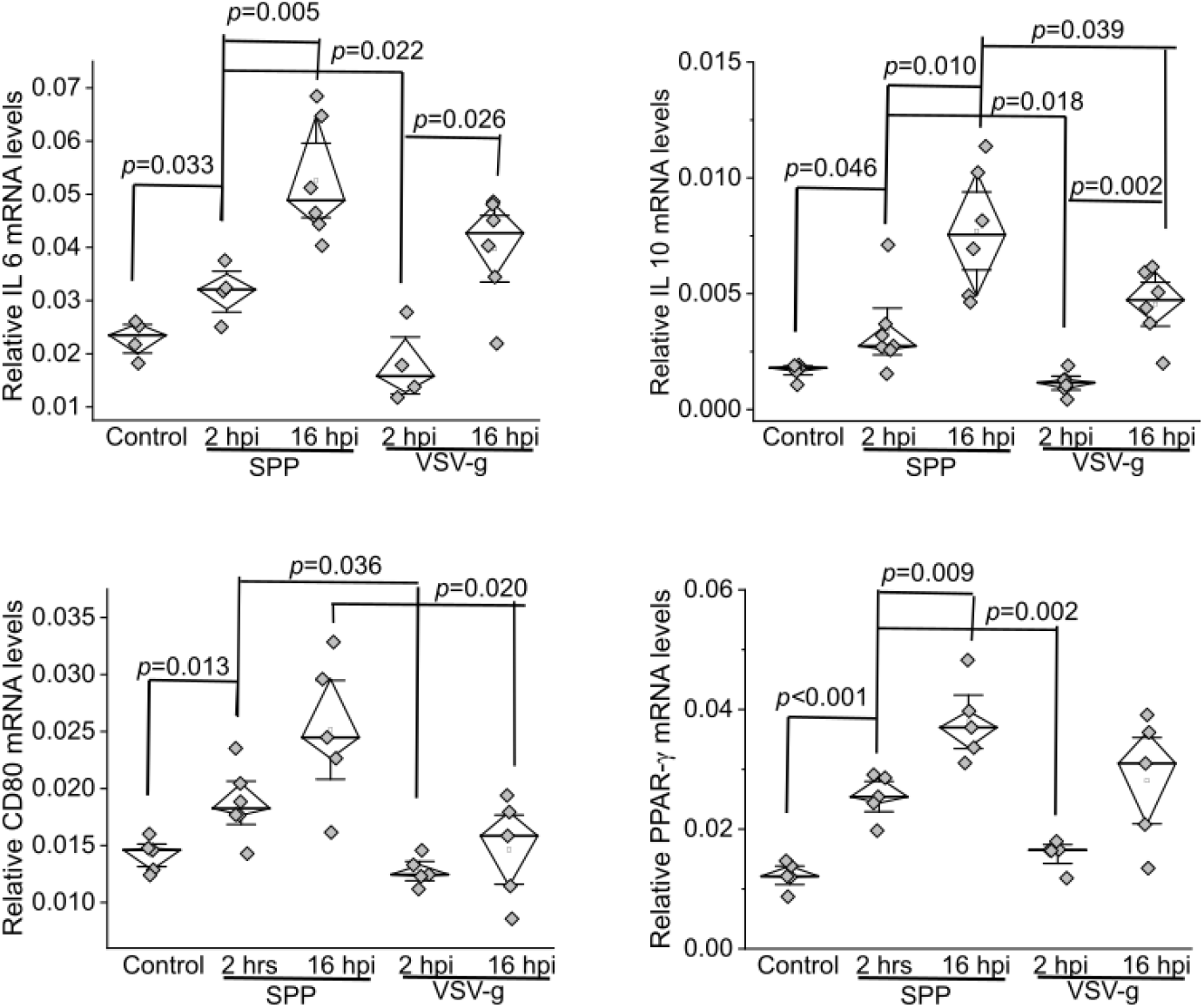
Upregulation of inflammatory factors in the RAW cells after being infected by Spike protein-pseudotyped (Spp) lentivirus or lentivirus carrying vesicular stomatitis virus glycoprotein (VSV-G) at 2 or 16 hours post-infection (hpi).

We next electroporated RAW cells with pcDNA-Spike expression plasmid or two control plasmid, pcDNA and EGFP-N2. The expression of EGFP two days post-electroporation showed 30%~40% transfection efficiency in RAW cells (Fig 6). The mRNA levels of IL 6, IL10, CD80 and PPAR-γ were significantly elevated in S protein-expression group than pcDNA or EGFP-N2 group (Fig 6).

**Fig 6.**
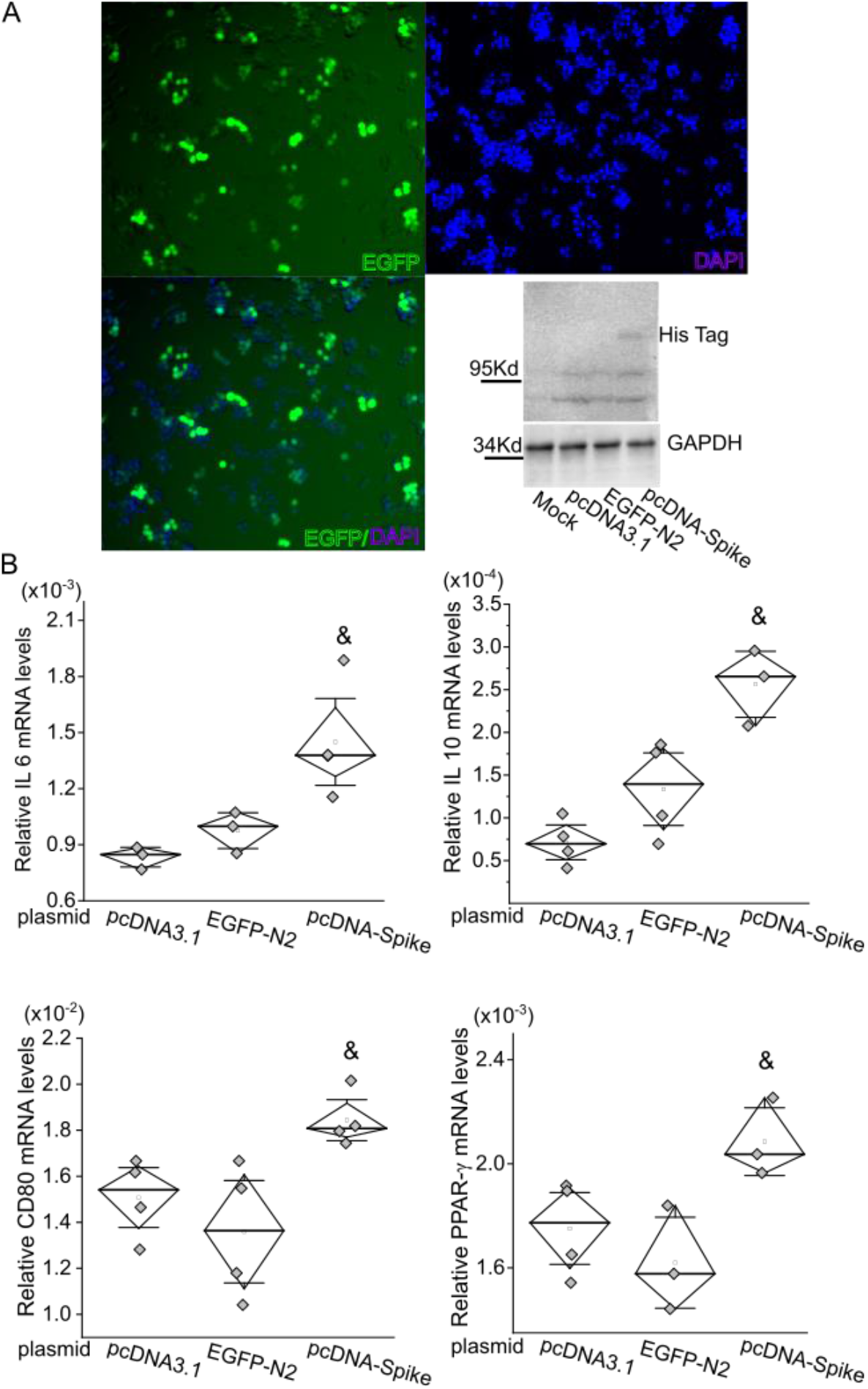
S protein of SARS-CoV-2 only induces upregulation of inflammatory factors in the RAW cells. (A) The expression of EGFP after electroporation of targeted plasmid into RAW cells to estimate the transfection efficiency. The expression of S protein in RAW is detected using an anti-His antibody by Western Blot. (B) The mRNA levels of IL-6, IL-10, CD80 and PPAR-γ are significantly upregulated after a forced expression of S protein. &, *p*<0.05 as comparison with control groups with electroporation of pcDNA3.1 blank vector or EGFP-N2 plasmid.

## Discussion

There is an urgent priority for global healthcare and the research community to mitigate the current pandemic of COVID-19. Animal models that can replicate viral transmission and subsequent pathological development are critical for this effort to understand the mechanisms of COVID-19 pathogenesis and develop effective anti-viral countermeasures. However, handling of specimens infected with SARS-CoV-2 requires high-security biosafety level 3 (BSL3) facilities and BSL3 work practices. We have developed the Spp lentivirus, which confers the Spike protein on the viral surface to investigate host tropism in a BSL2 setting. More importantly, using this virus, we have demonstrated the pathogenicity of the S protein in isolated cells and an animal model. Therefore, our data have proven that the Spp lentivirus, though it cannot completely replicate the infectious pathway and pathological process in human, is a very useful system to specifically investigate the S protein-mediated cell type susceptibility, host tropism for infection and pathogenicity.

SARS-CoV-2 RNA virus can be detected in swab samples from upper respiratory track from more than 80% of affected patients [16–18]. Viral RNAs are rarely detectable in blood of asymptomatic patients or non-hospitalized patients [16], but have been found more frequently in 10.5% ~ 67% hospitalized patients with severe symptoms [17–19]. Using a supersensitive RT-PCR RUO assay (low limit of detection at 625 copies of SARS-CoV-2 RNA/ml), blood SARS-CoV-2 has been detected in 53% in mild-to-moderate patients and 88% in critically ill patients, with levels of virus being associated with disease severity [20]. Viral shedding time (from positive to negative) of blood is shorter than that of nasal swab [18]. An animal study also has shown that intranasal inoculation of SARS-CoV-2 in a transgenic mouse model expressing human ACE2 results in high levels of viral infection in lungs with spread to other organs [21]. These reports demonstrate that SARS-CoV-2 can be disseminated from the respiratory system into the circulatory system and to many other vital organs during the late stage of disease, which will become one crucial determinant for a patient to develop severe symptoms. Therefore, it will be necessary to recapitulate how the disease progresses under this viremia condition in an animal model. In this study, we intravenously administrated Spp lentivirus into mice and attempted to develop an animal model to mimic part of these critical conditions from patients. Our data show high preference of Spp lentivirus residing in lung tissues and induction of an acute pneumonia, which is thought to be mediated or contributed by the S protein. Therefore, although it may not fully simulate the nature of the entry pathway of SARS-CoV-2 through upper airway to the lungs at the initial stage of the disease, this study has shown valuable information relevant to pathological progression of patients with viremia.

The S protein is thought to be the crucial glycoprotein on the surface of SARS-CoV-2 to mediate the viral entry of host cells. It can bind to ACE2 in the host cells, followed by an aid of TMPRSS2 cleavage, leading to endocytosis of viral/receptor complex into the host cells. ACE2 is present in type II alveolar cells, macrophages and endothelial cells in the lungs, which makes them targeted by SARS-CoV-2. Indeed, SARS-CoV-2 has been found in pneumocytes and endothelial cells from autopsy studies of COVID-19 patients [22]. In this study, we showed that Spp lentivirus was mainly present in LDLr^+^ type II alveolar cells and macrophages in the lungs. Those cells also express ACE2 [23, 24]. As ACE2 receptors in rodents have shown a much lower affinity to the SARS Spike protein as compared with human ACE2 receptor [21, 25, 26], the uptake of Spp lentivirus is less likely solely through the endogenous ACE2 in the lungs of mice. We speculate that other receptors or co-factors may facilitate the S protein-mediated viral entry of host cells via an ACE2-dependent or independent pathway for example NRP1-mediated entry pathway [8–10]. These data provide insight into the possibility of multiple factors/pathways involving SARS-CoV-2 entry of various target cells. The Spp lentivirus will allow us to investigate the mechanisms of viral entry into target cells in further detail in future studies.

Since the Spp is a replication-deficient virus, it will be unable to generate a full spectrum of pathological changes in host cells. We did not observe any evident lung inflammatory responses in mice at 2 hpi from either Spp or VSV-g. Mice at 24 hpi after Spp but not VSV-g administration developed acute, diffuse and evident lung inflammation. The pneumonia was transient and resolved 7 days after Spp administration (data not shown). This study demonstrate that Spp is able to replicate a portion of the acute lung pathology that SARS-CoV-2 causes in human. SARS-CoV-2-induced hyper-inflammation in the lungs is considered to cause disease progression. For example, the acute and diffuse lung injuries are supported by the evidence of upregulation of a series of serum cancer biomarkers in patients [27]. Our data show that a portion of M1 but not M2 macrophages exhibit rapid uptake of Spp lentivirus in the lungs (by 2 hpi), which provides direct evidence to support this notion. The severity of acute lung inflammation induced in our mouse model is more prominent in Spp than VSV-g. Furthermore, a panel of inflammatory factors are upregulated in the lungs post Spp infection, which also exhibit different patterns in the lungs with VSV-g infection. In consideration of the S protein being the only difference between Spp and VSV-g lentiviruses, these data lead us to speculate that the S protein may have executed a unique role in the development of this lung pathology. Indeed, over-expression of the S protein in the RAW cells can induce upregulation of the same panel of inflammatory factors, e.g. IL6, IL10, CD80 and PPAR-γ, demonstrating that the S protein has a function to induce intracellular pathological alterations. The detailed mechanisms underlying how the S protein contributes to the inflammatory reactions in macrophages are unknown and will be studied in future.

COVID-19 patients with metabolic-associated preconditions have a high risk to develop more severe symptoms. Our recent studies have shown that decreased levels of low and high density lipoprotein cholesterols are associated with severity and mortality of the COVID-19 [12, 13], which have been confirmed by many other reports [28–31]. The etiology of lipid abnormality in COVID-19 is likely multi-factorial including liver dysfunction, cytokine storm-induced alterations of lipid metabolism, virus-induced aberrant modulations in cholesterol synthesis and increased free radical signaling to facilitate modification and degradation of LDL-c [32, 33]. The S protein also has a binding pocket for fatty acids, steroids and cholesterol to modulate the host cell entrance [34, 35]. In this study, our data show that LDLr^+^ cells have the highest level of Spp lentivirus uptake in the lungs. This correlation together with previous reports indicates that lipid metabolism seems tightly associated with SARS-CoV-2 induced acute lung pathology. This data also suggests a possibility that LDLr may be directly involved in viral uptake, which shall be investigated in the future with the Spp lentivirus.

In conclusion, our data show that the Spp lentivirus can induce an acute and transient inflammatory response in the lungs of mice. The Spp lentivirus preferably targets those macrophages and pneumocytes with expression of LDLr in the lungs and lead to upregulation of IL6, IL10, CD80 and PPAR-γ. In addition, forced expression of the S protein can cause elevation of IL6, IL10, CD80 and PPAR-γ in RAW cells. Our results demonstrate that the S protein of SARS-CoV-2 can activate macrophages and contribute to induction of acute inflammations in the lungs.

## Author contributions

X.C. and Y.T. performed *in vivo* studies; V.N. performed *in vitro* RAW cell studies; Y.Z. generated Spp and VSV-G lentiviruses; C.G. and Y.R. participated experiments of cell culture, real time RT-PCR and immunoblot; W.C., D.F., H.A., and T.C. discussed the data and revised the manuscript; W.T. and T.C. covered the funding; W.T. designed and supervised the project.

## Acknowledgement

We greatly appreciate Dr. Mitzi Nagarkatti and Dr. Juhua Zhou from the Department of Pathology, Microbiology and Immunology at University of South Carolina School of Medicine for their very kind and generous help with real time PCR assays. We are also very thankful to the support and assistance from Instrumentation Resource Facility at University of South Carolina School of Medicine.

---

**Supplementary Table 1.**
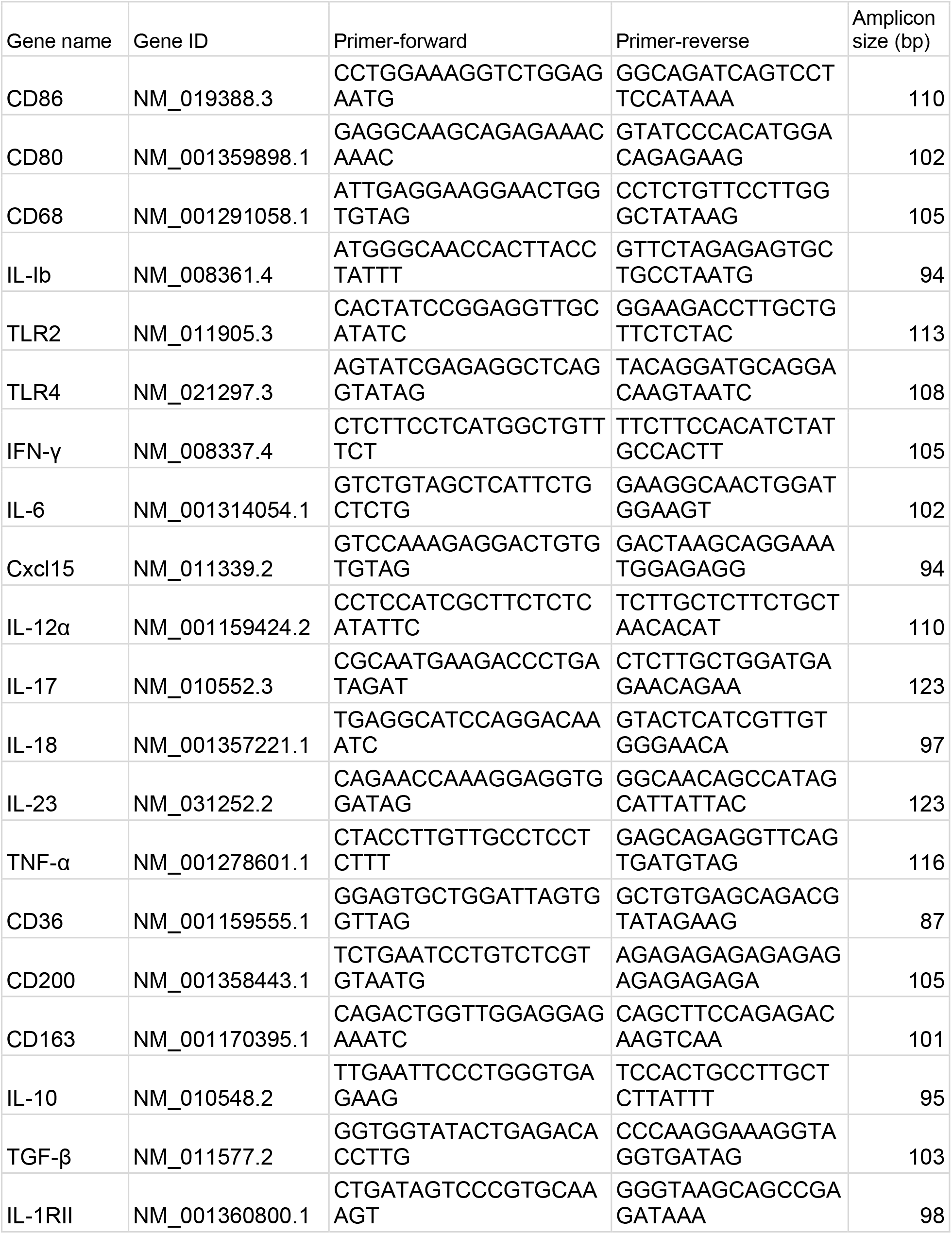
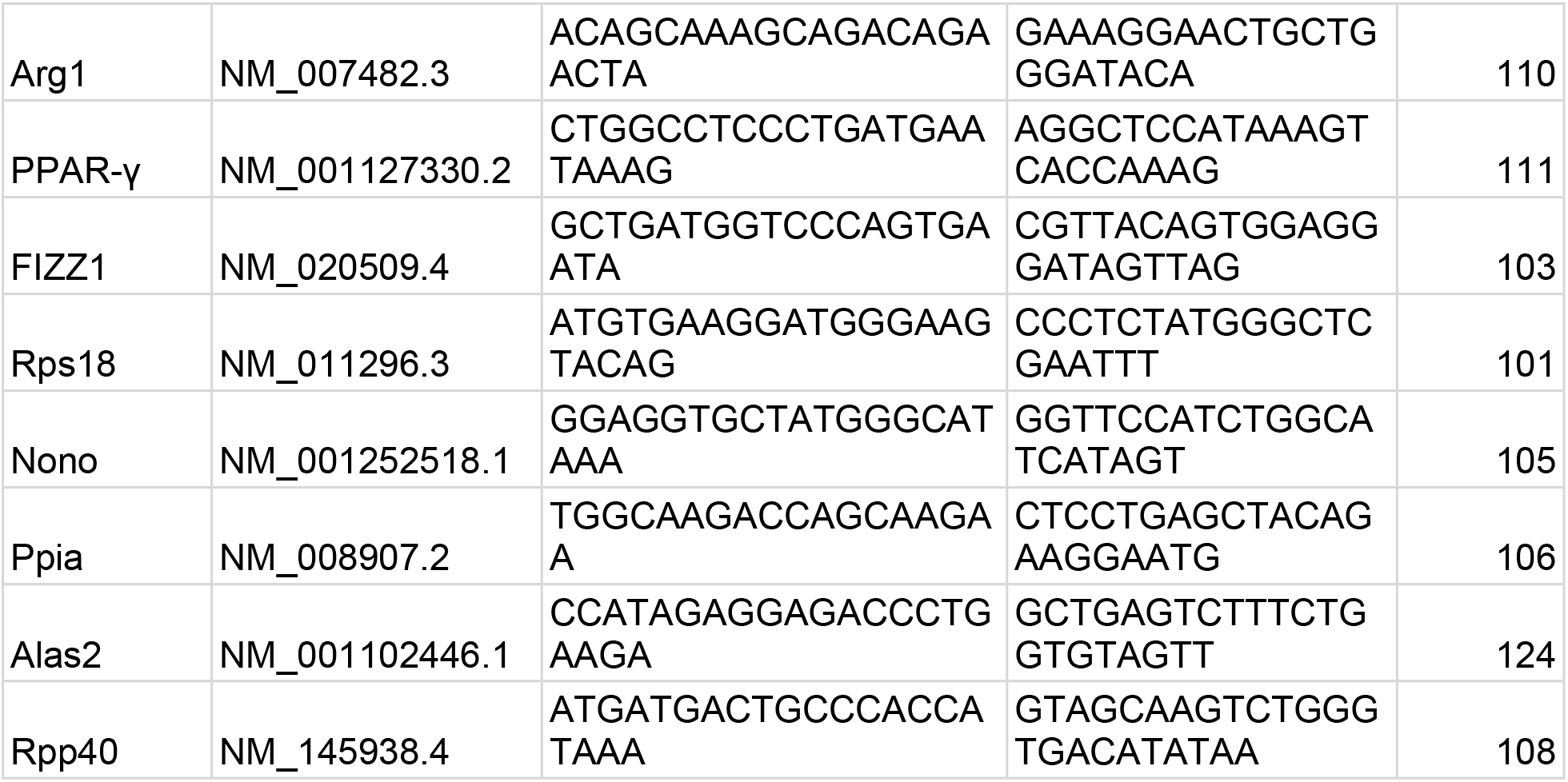
Primer lists for inflammatory genes

## References

1. CoronavirusResourceCenter. COVID-19 Dashboard by the Center for Systems Science and Engineering (CSSE) at Johns Hopkins University (JHU) 2020. Available from: https://coronavirus.jhu.edu/map.html.

2. The species Severe acute respiratory syndrome-related coronavirus: classifying 2019-nCoV and naming it SARS-CoV-2. Nat Microbiol. 2020. Epub 2020/03/04. doi: 10.1038/s41564-020-0695-z. PubMed PMID: 32123347.

3. Zhou P, Yang XL, Wang XG, Hu B, Zhang L, Zhang W, et al. A pneumonia outbreak associated with a new coronavirus of probable bat origin. Nature. 2020. Epub 2020/02/06. doi: 10.1038/s41586-020-2012-7. PubMed PMID: 32015507.

4. Walls AC, Park YJ, Tortorici MA, Wall A, McGuire AT, Veesler D. Structure, Function, and Antigenicity of the SARS-CoV-2 Spike Glycoprotein. Cell. 2020;181(2):281–92 e6. Epub 2020/03/11. doi: 10.1016/j.cell.2020.02.058. PubMed PMID: 32155444; PubMed Central PMCID: PMC7102599.

5. Hoffmann M, Kleine-Weber H, Pohlmann S. A Multibasic Cleavage Site in the Spike Protein of SARS-CoV-2 Is Essential for Infection of Human Lung Cells. Mol Cell. 2020;78(4):779–84 e5. Epub 2020/05/05. doi: 10.1016/j.molcel.2020.04.022. PubMed PMID: 32362314; PubMed Central PMCID: PMCPMC7194065.

6. Bestle D, Heindl MR, Limburg H, Van Lam van T, Pilgram O, Moulton H, et al. TMPRSS2 and furin are both essential for proteolytic activation of SARS-CoV-2 in human airway cells. Life Sci Alliance. 2020;3(9). Epub 2020/07/25. doi: 10.26508/lsa.202000786. PubMed PMID: 32703818; PubMed Central PMCID: PMCPMC7383062.

7. Ziegler C, Allon SJ, Nyquist SK, Mbano IM, Miao VN, Tzouanas CN, et al. SARS-CoV-2 receptor ACE2 is an interferon-stimulated gene in human airway epithelial cells and is enriched in specific cell subsets across tissues. Cell. 2020. doi: DOI: 10.1016/j.cell.2020.04.035.

8. Daly JL, Simonetti B, Klein K, Chen KE, Williamson MK, Anton-Plagaro C, et al. Neuropilin-1 is a host factor for SARS-CoV-2 infection. Science. 2020. Epub 2020/10/22. doi: 10.1126/science.abd3072. PubMed PMID: 33082294.

9. Cantuti-Castelvetri L, Ojha R, Pedro LD, Djannatian M, Franz J, Kuivanen S, et al. Neuropilin-1 facilitates SARS-CoV-2 cell entry and infectivity. Science. 2020. Epub 2020/10/22. doi: 10.1126/science.abd2985. PubMed PMID: 33082293.

10. Moutal A, Martin LF, Boinon L, Gomez K, Ran D, Zhou Y, et al. SARS-CoV-2 Spike protein co-opts VEGF-A/Neuropilin-1 receptor signaling to induce analgesia. Pain. 2020. Epub 2020/10/04. doi: 10.1097/j.pain.0000000000002097. PubMed PMID: 33009246.

11. Richardson S, Hirsch JS, Narasimhan M, Crawford JM, McGinn T, Davidson KW, et al. Presenting Characteristics, Comorbidities, and Outcomes Among 5700 Patients Hospitalized With COVID-19 in the New York City Area. Jama. 2020. Epub 2020/04/23. doi: 10.1001/jama.2020.6775. PubMed PMID: 32320003; PubMed Central PMCID: PMC7177629.

12. Fan J, Wang H, Ye G, Cao X, Xu X, Tan W, et al. Low-density lipoprotein is a potential predictor of poor prognosis in patients with coronavirus disease 2019. Metabolism. 2020:154243. Epub 2020/04/23. doi: 10.1016/j.metabol.2020.154243. PubMed PMID: 32320740; PubMed Central PMCID: PMC7166305.

13. Wei X, Zeng W, Su J, Wan H, Yu X, Cao X, et al. Hypolipidemia is associated with the severity of COVID-19. J Clin Lipidol. 2020. doi: doi: 10.1016/j.jacl.2020.04.008.

14. Voyno-Yasenetskaya TA, Dobbs LG, Erickson SK, Hamilton RL. Low density lipoprotein- and high density lipoprotein-mediated signal transduction and exocytosis in alveolar type II cells. Proc Natl Acad Sci U S A. 1993;90(9):4256–60. Epub 1993/05/01. doi: 10.1073/pnas.90.9.4256. PubMed PMID: 8483941; PubMed Central PMCID: PMCPMC46485.

15. Gowdy KM, Fessler MB. Emerging roles for cholesterol and lipoproteins in lung disease. Pulm Pharmacol Ther. 2013;26(4):430–7. Epub 2012/06/19. doi: 10.1016/j.pupt.2012.06.002. PubMed PMID: 22706330; PubMed Central PMCID: PMCPMC3466369.

16. Vm1 C, HF R, O A, D O, MB F, B K-S, et al. SARS-CoV-2 asymptomatic and symptomatic patients and risk for transfusion transmission. Transfusion. 2020. doi: 10.1111/trf.15841.

17. Chen W, Lan Y, Yuan X, Deng X, Li Y, Cai X, et al. Detectable 2019-nCoV viral RNA in blood is a strong indicator for the further clinical severity. Emerg Microbes Infect. 2020;9(1):5. doi: 10.1080/22221751.2020.1732837.

18. Fang Z, Zhang Y, Hang C, Ai J, Li S, Zhang W. Comparisons of viral shedding time of SARS-CoV-2 of different samples in ICU and non-ICU patients. J Infect. 2020. Epub 2020/03/27. doi: 10.1016/j.jinf.2020.03.013. PubMed PMID: 32209381; PubMed Central PMCID: PMC7118636.

19. Buetti N, Patrier J, Le Hingrat Q, Loiodice A, Bouadma L, Visseaux B, et al. Risk factors for SARS-CoV-2 detection in blood of critically ill patients. Clin Infect Dis. 2020. Epub 2020/09/03. doi: 10.1093/cid/ciaa1315. PubMed PMID: 32875309; PubMed Central PMCID: PMC7499490.

20. Veyer D, Kerneis S, Poulet G, Wack M, Robillard N, Taly V, et al. Highly sensitive quantification of plasma SARS-CoV-2 RNA shelds light on its potential clinical value. Clin Infect Dis. 2020. Epub 2020/08/18. doi: 10.1093/cid/ciaa1196. PubMed PMID: 32803231; PubMed Central PMCID: PMC7454373.

21. Winkler ES, Bailey AL, Kafai NM, Nair S, McCune BT, Yu J, et al. SARS-CoV-2 infection of human ACE2-transgenic mice causes severe lung inflammation and impaired function. Nat Immunol. 2020;21(11):1327–35. Epub 2020/08/26. doi: 10.1038/s41590-020-0778-2. PubMed PMID: 32839612; PubMed Central PMCID: PMCPMC7578095.

22. Fox SE, Akmatbekov A, Harbert JL, Li G, Quincy Brown J, Vander Heide RS. Pulmonary and cardiac pathology in African American patients with COVID-19: an autopsy series from New Orleans. Lancet Respir Med. 2020;8(7):681–6. Epub 2020/05/31. doi: 10.1016/S2213-2600(20)30243-5. PubMed PMID: 32473124; PubMed Central PMCID: PMCPMC7255143.

23. Wiener RS, Cao YX, Hinds A, Ramirez MI, Williams MC. Angiotensin converting enzyme 2 is primarily epithelial and is developmentally regulated in the mouse lung. J Cell Biochem. 2007;101(5):1278–91. Epub 2007/03/07. doi: 10.1002/jcb.21248. PubMed PMID: 17340620; PubMed Central PMCID: PMCPMC7166549.

24. Keidar S, Gamliel-Lazarovich A, Kaplan M, Pavlotzky E, Hamoud S, Hayek T, et al. Mineralocorticoid receptor blocker increases angiotensin-converting enzyme 2 activity in congestive heart failure patients. Circ Res. 2005;97(9):946–53. Epub 2005/09/24. doi: 10.1161/01.RES.0000187500.24964.7A. PubMed PMID: 16179584.

25. Li W, Greenough TC, Moore MJ, Vasilieva N, Somasundaran M, Sullivan JL, et al. Efficient replication of severe acute respiratory syndrome coronavirus in mouse cells is limited by murine angiotensin-converting enzyme 2. J Virol. 2004;78(20):11429–33. Epub 2004/09/29. doi: 10.1128/JVI.78.20.11429-11433.2004. PubMed PMID: 15452268; PubMed Central PMCID: PMCPMC521845.

26. Letko M, Marzi A, Munster V. Functional assessment of cell entry and receptor usage for SARS-CoV-2 and other lineage B betacoronaviruses. Nat Microbiol. 2020;5(4):562–9. Epub 2020/02/26. doi: 10.1038/s41564-020-0688-y. PubMed PMID: 32094589; PubMed Central PMCID: PMCPMC7095430.

27. Wei X, Su J, Yang K, Wei J, Wan H, Cao X, et al. Elevations of serum cancer biomarkers correlate with severity of COVID-19. J Med Virol. 2020. Epub 2020/04/30. doi: 10.1002/jmv.25957. PubMed PMID: 32347972; PubMed Central PMCID: PMCPMC7267262.

28. Hu X, Chen D, Wu L, He G, Ye W. Declined serum high density lipoprotein cholesterol is associated with the severity of COVID-19 infection. Clin Chim Acta. 2020. Epub 2020/07/13. doi: 10.1016/j.cca.2020.07.015. PubMed PMID: 32653486; PubMed Central PMCID: PMC7350883.

29. Sorokin AV, Karathanasis SK, Yang ZH, Freeman L, Kotani K, Remaley AT. COVID-19-Associated dyslipidemia: Implications for mechanism of impaired resolution and novel therapeutic approaches. Faseb J. 2020. Epub 2020/06/27. doi: 10.1096/fj.202001451. PubMed PMID: 32588493; PubMed Central PMCID: PMC7361619.

30. Wang G, Zhang Q, Zhao X, Dong H, Wu C, Wu F, et al. Low high-density lipoprotein level is correlated with the severity of COVID-19 patients: an observational study. Lipids Health Dis. 2020;19(1):204. Epub 2020/09/08. doi: 10.1186/s12944-020-01382-9. PubMed PMID: 32892746; PubMed Central PMCID: PMCPMC7475024.

31. Tanaka S, De Tymowski C, Assadi M, Zappella N, Jean-Baptiste S, Robert T, et al. Lipoprotein concentrations over time in the intensive care unit COVID-19 patients: Results from the ApoCOVID study. PLoS One. 2020;15(9):e0239573. Epub 2020/09/25. doi: 10.1371/journal.pone.0239573. PubMed PMID: 32970772; PubMed Central PMCID: PMCPMC7514065.

32. Bojkova D, Klann K, Koch B, Widera M, Krause D, Ciesek S, et al. Proteomics of SARS-CoV-2-infected host cells reveals therapy targets. Nature. 2020. Epub 2020/05/15. doi: 10.1038/s41586-020-2332-7. PubMed PMID: 32408336.

33. Cao X, Yin R, Albrecht H, Fan D, Tan W. Cholesterol: A new game player accelerating vasculopathy caused by SARS-CoV-2? Am J Physiol Endocrinol Metab. 2020;319(1):E197–E202. Epub 2020/06/06. doi: 10.1152/ajpendo.00255.2020. PubMed PMID: 32501731; PubMed Central PMCID: PMCPMC7347957.

34. Toelzer C, Gupta K, Yadav SKN, Borucu U, Davidson AD, Kavanagh Williamson M, et al. Free fatty acid binding pocket in the locked structure of SARS-CoV-2 spike protein. Science. 2020;370(6517):725–30. Epub 2020/09/23. doi: 10.1126/science.abd3255. PubMed PMID: 32958580.

35. Shoemark D, Colenso C, Toelzer C, Gupta K, Sessions R, Davidson A, et al. Molecular Simulations suggest Vitamins, Retinoids and Steroids as Ligands binding the Free Fatty Acid Pocket of SARS-CoV-2 Spike Protein. Chemrxiv. 2020. doi: https://doi.org/10.26434/chemrxiv.13143761.v1

